# Reconstruction of Cellular Trajectories with Biologically Interpretable States

**DOI:** 10.64898/2026.11.03.763902

**Authors:** Yuhan Wang, Magnus Rattray, David Barber

## Abstract

Elucidating the trajectories of cellular evolution provides biologists with deeper insights into biological systems and disease progression, ultimately facilitating more informed discovery of mechanisms and therapeutic design. Single-cell RNA sequencing (scRNA-seq) is widely used to study cellular evolution. However, it faces two primary limitations. First, because the process destroys cells during gene-expression measurement, it is impossible to track the gene-expression profile of the same cell throughout the entire trajectory. Second, because scRNA-seq captures cells at discrete time points, it provides only static snapshots rather than continuous trajectories, limiting its ability to resolve the full dynamics of cellular transitions. Consequently, various computational approaches have been developed to address these challenges. However, existing models fail to capture the hidden state of each cell, which is essential for reconstructing gene expression at unobserved time points. As the gene expression profiles depend on the cell state, and may vary according to time, we propose a Latent Cell-State Model, where we approximate cell state transition as a Markov chain and obtain the gene expression of each cell at unobserved timepoints depending on the cell state, time, and a latent variable that represents the high-dimensional gene expression profile of a cell. Our model is able to infer biologically meaningful cell states and generate high-quality gene expression profiles at unobserved timepoints, showing potential for accurate trajectory reconstruction.

## 1 Introduction

Cell evolution is inherently dynamic, with molecular and cellular components continuously changing over time. To capture these processes, single-cell gene profiling methods are widely used to investigate cellular differentiation, as well as to better understand drug responses and disease progression. However, single-cell RNA-sequencing (scRNA-seq) is not able to fully capture the temporal dynamics of cell states [1]. Firstly, it cannot profile the same cell across multiple time points, as the measurement process destroys the cell, yielding only a snapshot of its state at a single time. Secondly, the sampling is often sparse, leading to information loss between consecutive time points. As a result, rapid changes occurring during unobserved intervals are difficult to capture [2]. To better characterize these challenges and define the problem, we illustrate it in Fig. 1(a). A total number of cell samples *S* are observed at pre-selected time points *T* from time 0 to time *T*. For each cell sample *s ∈* {1, …, *S*}, we have a collection of *G* genes. Each gene *g ∈* {1, …, *G*} has a discrete expression value 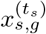, at timepoint *t*_*s*_ *∈ T* when the cell is destroyed to measure the gene expression.

**Fig. 1:**
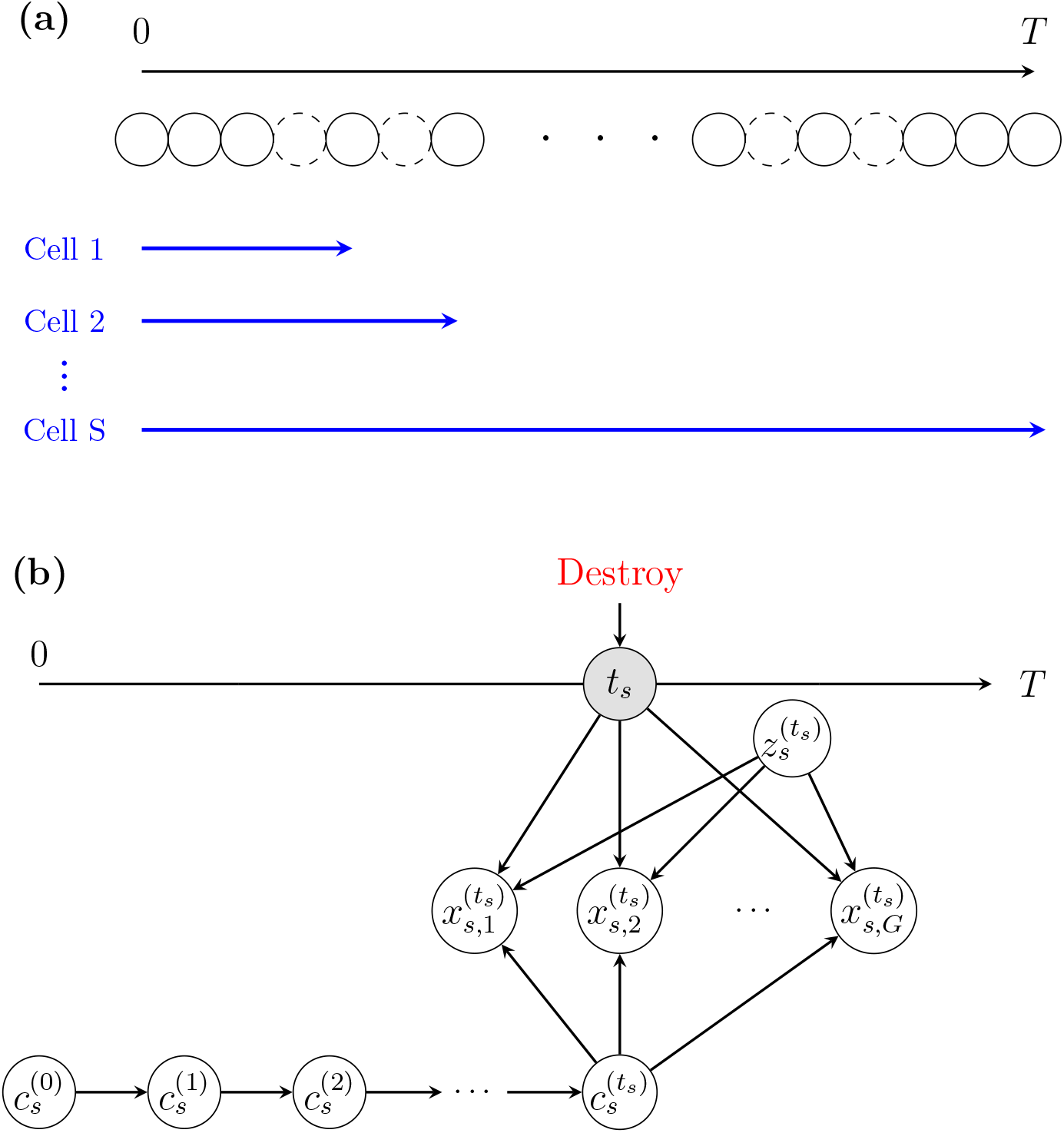
**(a)** scRNA-seq captures gene-expression for each cell sample *s*. For each cell sample *s ∈* {1, …, *S*}, the cellular evolution starts at time 0 and continues until time *t*_*s*_ marked by the blue arrowhead when the cell is destroyed and the gene expression measured. The dashed circles represent unobserved timpoints, whereas the solid circles mark observed timepoints where scRNA-seq data are captured. **(b)** Latent Cell-State Model: For one observed cell sample *s*, where *s ∈* {1, …, *S*}, we have a collection of *G* genes. Each gene *g ∈* {1, …, *G*} has a discrete expression value 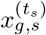, at timepoint *t* when the cell is destroyed for gene expression measurement. We assume that the cell starts to evolve at time 0 and transitions according to a Markov chain and becomes 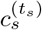 at time *t*_*s*_, when the cell is destroyed for measurement. Then, the expression of each gene 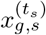 at time *t*_*s*_ can be obtained based on current time *t*_*s*_, latent variable 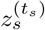 and current cell state 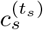.

Our interest is to recover the gene expression profiles of cells at unobserved times *t* ∉ *T* based on the gene expression profiles of all cells observed along the evolution trajectory *X*. To tackle this problem, we assume that gene expression is driven by a discrete unobserved (latent) cell-state *c*, representing which of *C* possible clusters the gene expression belongs to, and is also affected by the time *t* at which the cell is destroyed for measurement. We assume that each cell evolves from some unknown initial state and transitions to later states until a pre-specified time when the cell is destroyed and the gene expressions measured [3].

Specifically, as shown in Fig. 1(b), each cell sample *s ∈* {1, …, *S*}, evolves from an initial state 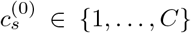 according to a Markov transition matrix *P* that ensures *c*^*t*+1^ is at least as large as *c*^*t*^, until time *t*_*s*_ when the cell is destroyed in order to measure its gene expression. Moreover, alongside cell state 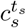, time *t*_*s*_ directly influences the gene expression to capture continuous changes in expression through time. To facilitate modelling dependencies in the gene expression values for each cell, we use a low-dimensional continuous latent space *z* shared across all cell samples, cell-states, and times^1^.

Our resulting Latent Cell-State Model is given by

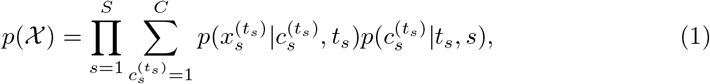

where

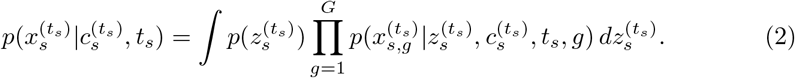

The cell-state distribution is given by the marginal of a Markov chain,

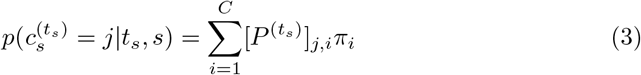

where *P* is a constrained transition matrix to ensure transitions to lower states are not allowed and *π* is the initial state distribution.

Details of the model and implementation are provided in Section 4, including how the expression profile is generated from the underlying low-dimensional representation, cell-state, and the unobserved timepoint of interest. For our model, the resulting likelihood *p*(*X*) is computationally intractable and we use a variational approach to form a lower bound on log *p*(*X*). All parameters of the model are then learned by maximising this lower bound approximation to the overall likelihood. In addition to allowing us to then predict expression values for unobserved timepoints, our model allows us to cluster gene expression into a small number of cell-states *C*, enabling us to interpret complex high-dimensional gene expression through a small number of macro cell-states.

## 2 Results

### 2.1 Benchmarking Temporal Interpolation

We compare the performance of our model against three state-of-the-art approaches scNode[4], scDiffusion [5] and ARTEMIS [6] on the Pancreatic dataset [7] and the Waddington-OT dataset [8]. The datasets are detailed in Section 4.5. For quantitative evaluation, we evaluate the discrepancy between generated and real test samples using maximum mean discrepancy (MMD) [9], which is detailed in Supplementary Note 3. For qualitative evaluation, we use uniform manifold approximation and projection (UMAP) [10] to visualize the quality of the cell samples generated by different approaches, which is also detailed in Supplementary Note 3.

#### 2.1.1 Interpolation on Pancreatic Development

In the Pancreatic dataset [7], we focus on stage 5, which covers the period from day 0 to day 7 (8 timepoints) and represents the critical transition where pancreatic progenitors undergo an epithelial-mesenchymal-like transition to delaminate and commit to endocrine fate. To assess the ability of the model to predict gene expression at unobserved timepoints, we hold out day 3 and day 6 from the Pancreatic dataset, where day 3 represents an intermediate stage and day 6 corresponds to a later internal stage of differentiation. To compare the performance of our and competing models, 2000 samples are generated for each held-out day from each model.

As shown in Fig. 2(a), the MMD between real samples and the samples generated by our model is 0.011 for day 3 and 0.006 for day 6, showing that our method out-performs the three comparison methods on both days. Moreover, the two dimensional UMAP on the generated and real cell samples at held-out timepoints in Fig. 2(b) shows that our model can capture a distribution that closely resembles the real one for both day 3 and day 6. In contrast, the comparison methods struggle to generate samples that fully cover the regions occupied by the real data.

**Fig. 2:**
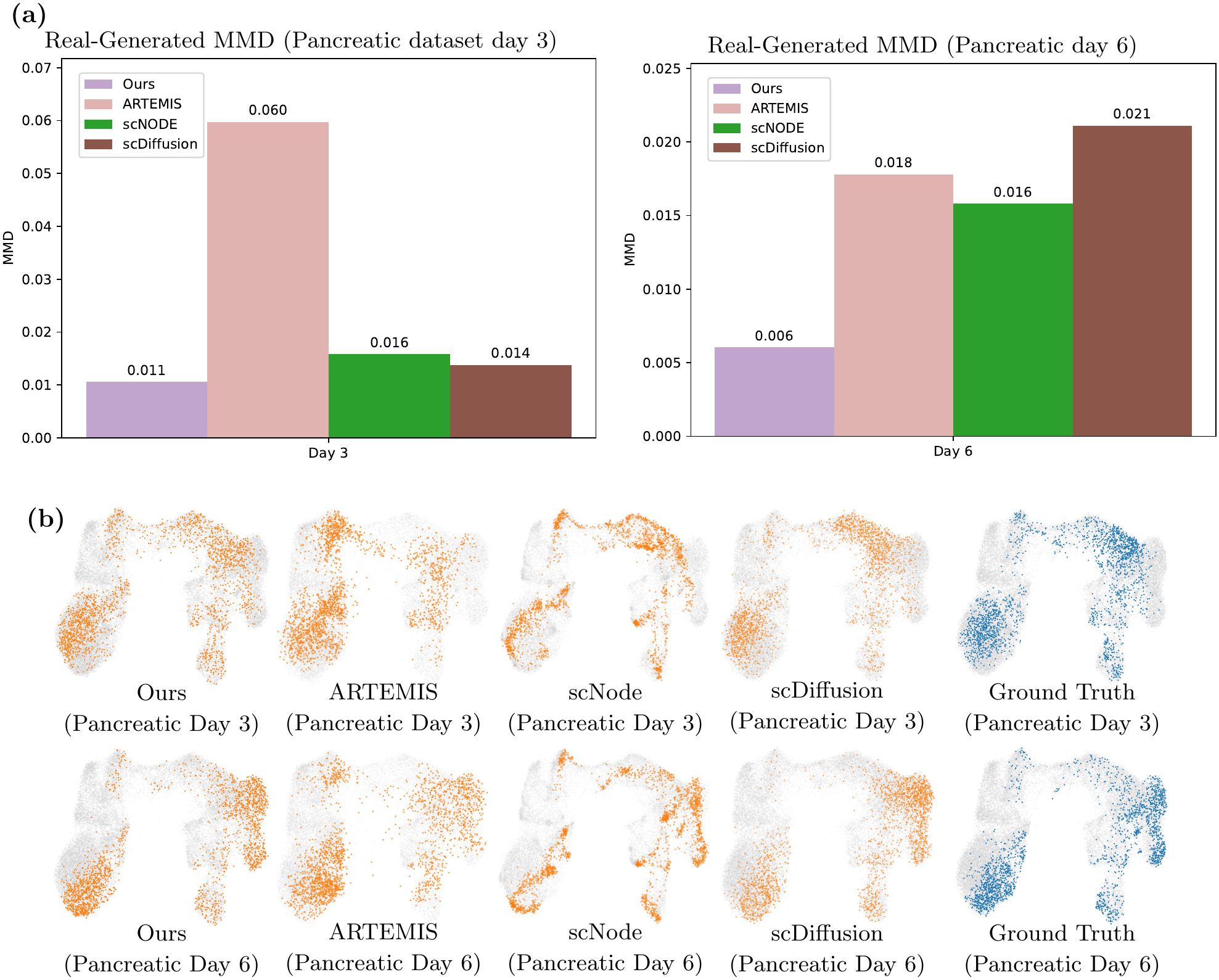
(a) MMD between real and generated samples at days 3 and 6 across different methods in the Pancreatic dataset. Different colours denote different methods. (b) 2D-UMAP visualization comparing generated and real samples from different methods at days 3 and 6 in the Pancreatic dataset. Generated samples at the held-out dates are highlighted in orange, real samples at the held-out dates are highlighted in dark blue, and real samples from non-held-out dates are shown in gray. The first row shows the 2D-UMAP visualization comparing real and generated samples at day 3, while the second row shows the corresponding comparison at day 6.

#### 2.1.2 Interpolation on Reprogramming Dynamics

In the Waddington-OT dataset [8], we focus on data from day 0 to day 8, sampled at half-day intervals (17 timepoints), which comprise mouse embryonic fibroblasts undergoing reprogramming into induced pluripotent stem cells (iPSCs). We use days 3.5 and 4 as hold-out timepoints to test interpolation performance. 1000 samples are generated for each held-out day from our model and the comparison methods. The generative performances are illustrated in Fig. 3(a). As shown in Fig. 3(a), the MMD between the real cell samples and the samples generated by our model is 0.030 for day 3.5 and 0.018 for day 4, both of which are significantly lower than those of the three comparison methods. Additionally, the first row of Fig. 3(b) shows the two-dimensional UMAP embedding of the generated and real samples on day 3.5. It can be observed that, under our model, the generated samples align closely with the real data distribution in the UMAP embedding space. For the comparison methods, although the generated samples partially overlap with the real samples, they fail to fully cover the region occupied by the real data, resulting in an incomplete representation of the true distribution. For the two-dimensional UMAP embedding of the generated and real samples on day 4, as shown in the second row of Fig. 3(b), a consistent pattern is observed.

**Fig. 3:**
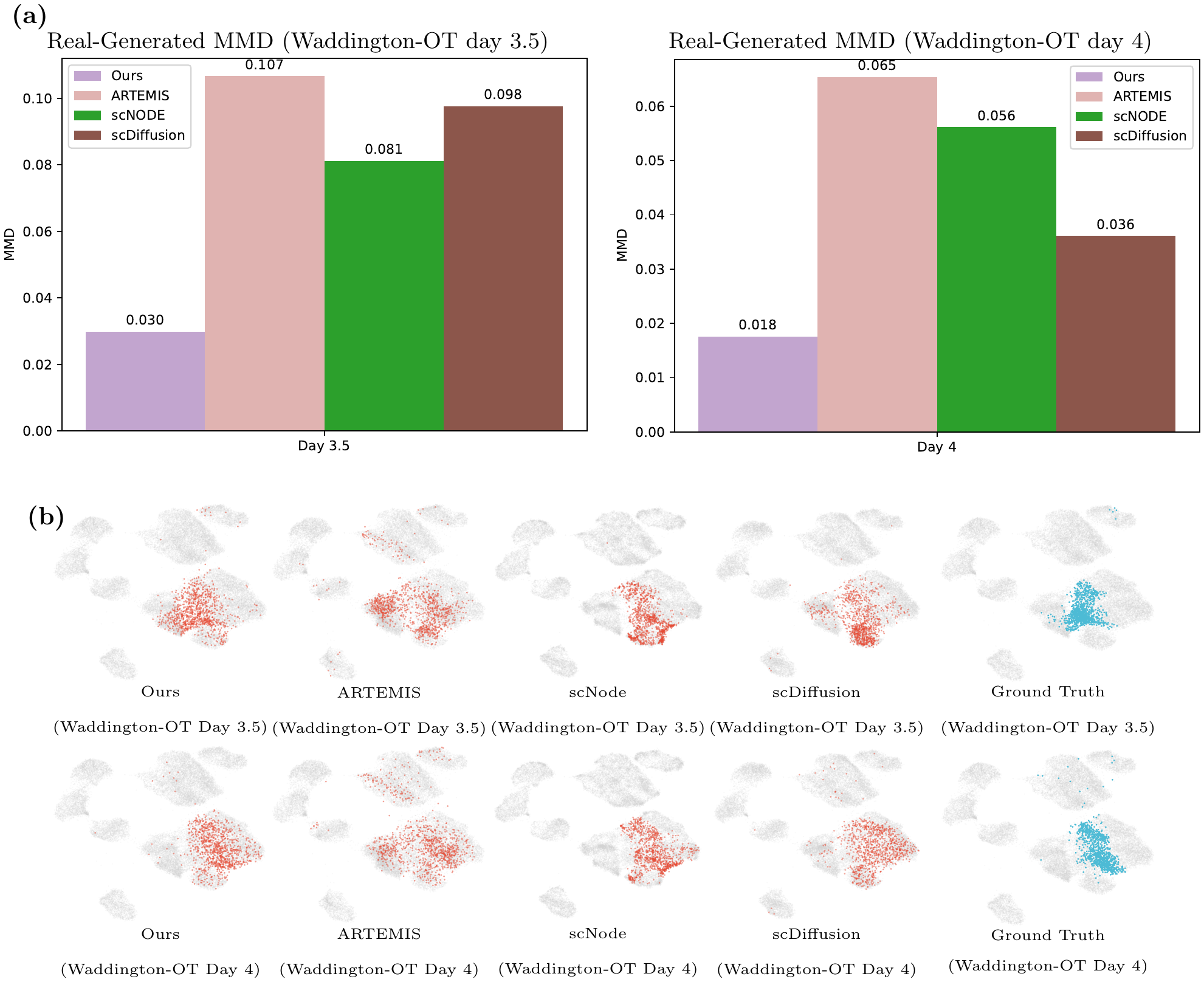
(a) MMD between real and generated samples at days 3.5 and 4 across different methods in the Waddington-OT dataset. Different colours denote different methods. (b) 2D-UMAP visualization comparing generated and real samples from different methods at days 3.5 and 4 in the Waddington-OT dataset. Generated samples at the held-out dates are highlighted in red, real samples at the held-out dates are highlighted in light blue, and real samples from non-held-out dates are shown in gray. The first row shows the 2D-UMAP visualization comparing real and generated samples at day 3.5, while the second row shows the corresponding comparison at day 4.

### 2.2 Biological Characterization of Model-Identified Cell States

A key feature of the Latent Cell-State Model is that gene expression is driven by a small number of cell-states *C* as the cell evolves. We aim to capture the cell-state using all cell samples at observed timepoints and then generate gene expression with the corresponding cell-state along the entire trajectory. Our interest here is to investigate the biological significance of the learnt cell-states.

#### 2.2.1 Determining Endocrine Identities

We first focus on the Pancreatic dataset, where the transition from pancreatic progenitors toward endocrine lineages is captured, including *β*-like, *α*-like, and SC-EC-like cells with a separate branch that follows a non-endocrine fate. To evaluate the cell states generated by our methods, we generate two thousands samples per timepoint along the trajectory, using the trained model. To understand the cell states generated by our model, we visualize these cell samples using the 2-dimensional UMAP, where cell samples are coloured by their corresponding generated cell state numbers, as shown in Fig. 4(a). To characterize each cell state, we identify its top 20 differentially expressed genes using the Wilcoxon rank-sum test [11], with genes ranked by the test statistic. All top genes are illustrated in Supplementary Fig. 1. Then, the dendrogram shown in Fig. 4(b) is generated by hierarchical clustering of cell states based on the first 50 principal components (PCs) obtained from PCA [12] of the normalized and log-transformed gene expression data. To further explore the biological characteristics of each cell state, we visualize the mean expression of selected canonical marker genes among expressing cells alongside the percentage of expressing cells across all generated samples using matrix and dot plots [11], as shown in Figs. 4(d) and (c) (See Supplementary Note 4 for further details on the UMAP visualization procedure, the identification and selection of marker genes, and the methods used for hierarchical clustering and gene expression visualization).

**Fig. 4:**
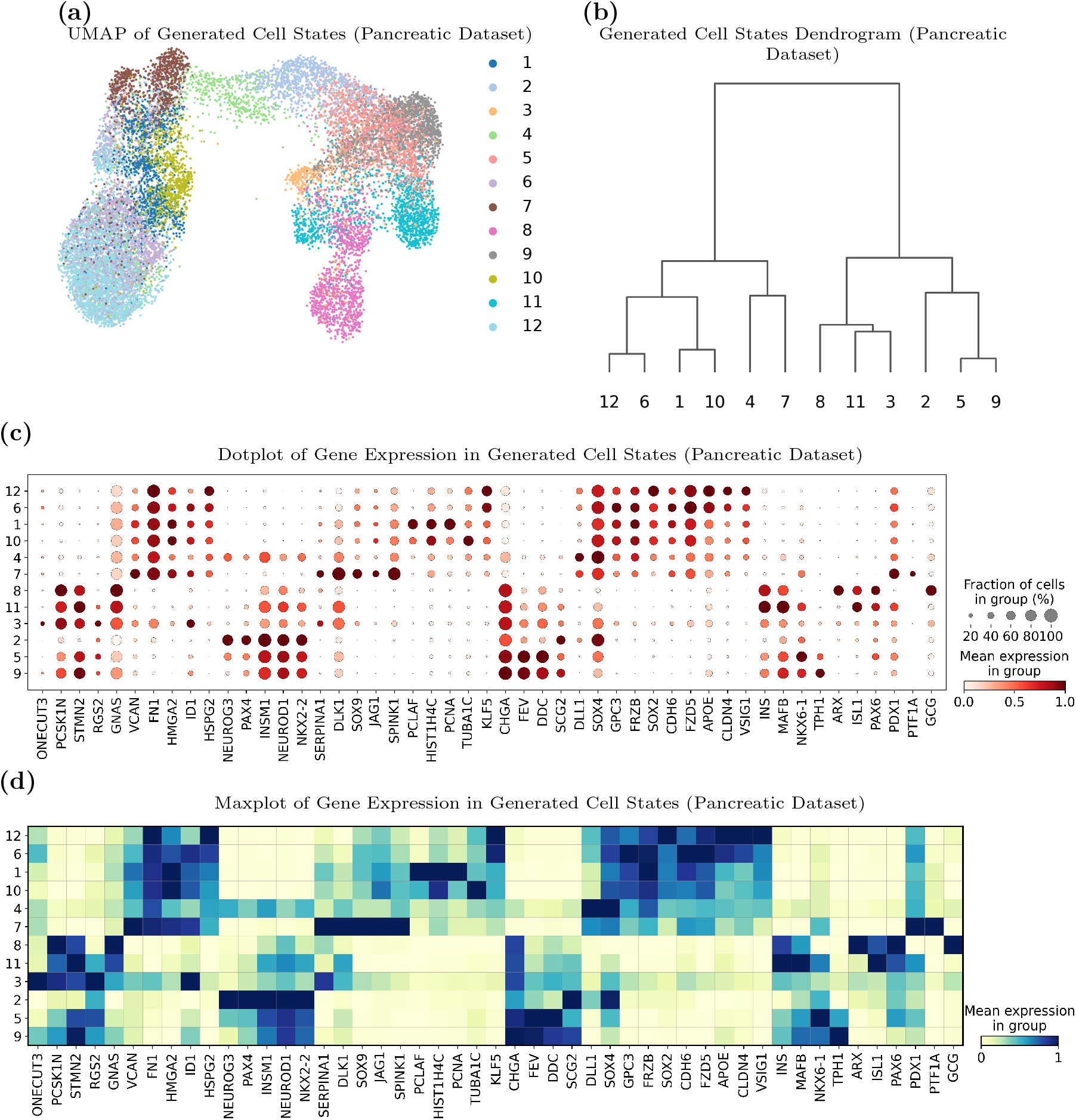
Pancreatic Dataset: (a) 2D-UMAP visualization of generated cell samples colored by generated cell state, where different colours indicates different cell states; (b) Dendrogram showing the hierarchical relationships among generated cell states. (c) Dot plot showing the mean expression of canonical marker genes across generated cell states. Dot colour indicates the mean expression level, and dot size represents the fraction of cells expressing each gene. (d) Matrix plot showing the mean expression of canonical marker genes across generated cell states, with colour gradients indicating differences in gene expression levels.

As shown in Fig. 4(b), hierarchical clustering reveals two major transcriptional branches, with one showing more prominent endocrine-associated expression patterns and the other encompassing a heterogeneous range of cellular programmes. We further examined the marker-expression patterns within these branches using the dot plot in Fig. 4(c) and the matrix plot in Fig. 4(d), and used these patterns to annotate the generated cell states. State 7 retains a pancreatic progenitor-like profile, characterized by prominent *SOX9* and *PDX1* expression [13, 14]. State 4 is consistent with an early endocrine-induction state, characterized by prominent *DLL1* expression together with emerging *NEUROG3*, with additional support from *SOX4* expression [15, 16]. State 2 corresponds to a more established endocrine-induction state, with prominent *NEUROG3* and *PAX4* expression, accompanied by *INSM1, NEUROD1*, and *NKX2-2* [7, 15, 17]. State 5 represents a differentiating endocrine intermediate, characterized by expression of *INSM1, NEUROD1*, and *NKX2-2*, together with increased expression of endocrine-associated genes including *CHGA, FEV*, and *DDC* [7, 18]. State 9 displays a strong *TPH1*-associated endocrine programme together with *CHGA, FEV*, and *DDC*, consistent with an SC-EC-like identity [7]. State 8 is characterized by prominent *ARX* and *GCG* expression, supporting an *α*-like identity, whereas State 11 shows prominent *INS* and *NKX6-1* expression with little *ARX* and*GCG* signal, consistent with a *β*-like endocrine state [7]. State 3 exhibits an immature endocrine-like profile, with *CHGA* expression but without the strong lineage-defining markers observed in States 8 and 11. Its expression of *ID1*, a developmental regulator associated with pancreatic progenitor populations, may indicate the retention of progenitor-associated transcriptional features [19]. States 1 and 10 represent closely related proliferative non-endocrine states but exhibit distinct proliferation-associated expression patterns. State 1 shows prominent expression of the replication-associated genes *HIST1H4C*, and *PCNA*, consistent with active DNA replication [20–22]. In contrast, State 10 exhibits a distinct cell-cycle-associated transcriptional programme, characterized by relatively strong expression of *HIST1H4C* and *TUBA1C* [20, 23]. States 6 and 12 exhibit closely related non-endocrine epithelial transcriptional profiles, characterized by shared expression of *FZD5* and *CLDN4* [19, 24]. State 12 displays relatively elevated *SOX2* expression, suggesting features of an earlier progenitor-like state [25], whereas State 6 exhibits a related progenitor-associated transcriptional profile with differences in the expression of developmental signalling-related genes, including *ID1* [19]. A complementary model-based analysis of the gene-expression distribution *p*(*X*_*g*_ | *c*) (defined in Supplementary Note 5), where *X*_*g*_ denotes the gene-expression variable and *g* indexes the canonical genes, showed broadly concordant state-level transcriptional programmes (Supplementary Fig. 5 (a)).

#### 2.2.2 Characterization of Reprogramming States

We then focus on the Waddington-OT dataset, which captures the emergence of stromal- and MET-associated transcriptional programs during early reprogramming. We generate one thousand samples per timepoint along the trajectory for evaluation. To understand the cell states generated by our model, we illustrate the two dimensional UMAP of all generated cell samples coloured by their corresponding cell state numbers, shown in Fig. 5(a). To characterize each cell state, we also present the top 20 highly expressed genes for each state (all selected genes are shown in Supplementary Fig. 2). Then, Fig. 5(b) is generated by hierarchical clustering of cell states based on the first 50 principal components (PCs) obtained from PCA [12] of the normalized and log-transformed gene expression data. To further explore the biological characteristics of each cell state, we visualize the mean expression of selected canonical marker genes among expressing cells alongside the percentage of expressing cells across all generated samples using matrix and dot plots defined in [11], as shown in Fig. 5(d) and Fig. 5(c) (See Supplementary Note 4 for further details on the UMAP visualization procedure, the identification and selection of marker genes, and the methods used for hierarchical clustering and gene expression visualization).

**Fig. 5:**
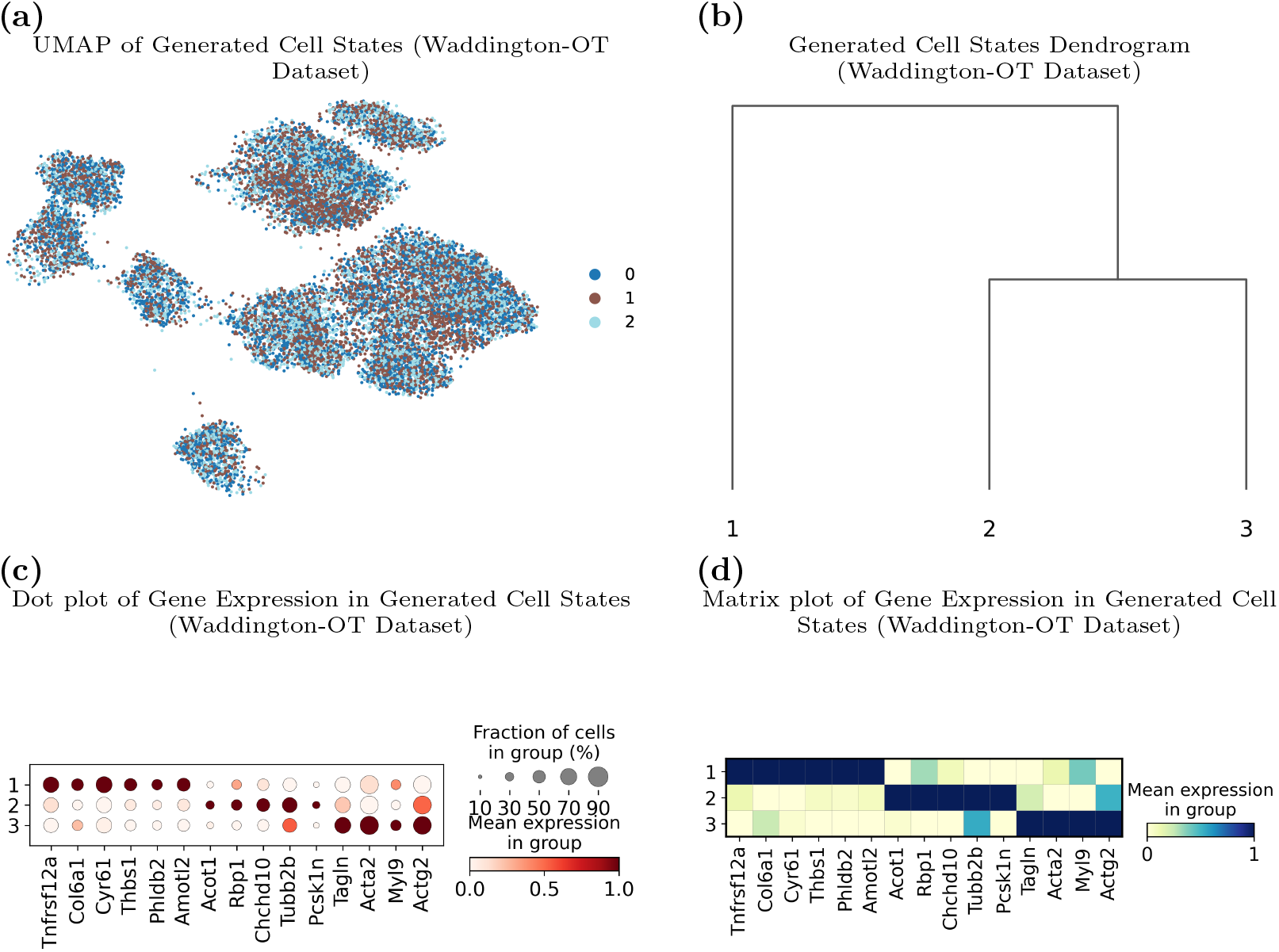
Waddington-OT Dataset: (a) 2D-UMAP visualization of generated cell samples colored by generated cell state, where different colours indicates different cell states; (b) Dendrogram showing the hierarchical relationships among generated cell states. (c) Dot plot showing the mean expression of canonical marker genes across generated cell states. Dot colour indicates the mean expression level, and dot size represents the fraction of cells expressing each gene. (d) Matrix plot showing the mean expression of canonical marker genes across generated cell states, with colour gradients indicating differences in gene expression levels.

The cell states were identified based on the gene markers described in [8]. The selected marker genes revealed distinct transcriptional profiles across the three generated cell states. State 1 preferentially expressed *Col6a1, Cyr61, Thbs1*, and *Tnfrsf12a*, indicating extracellular matrix- and stromal-associated features. State 2 exhibited a more heterogeneous transcriptional profile, characterized by intermediate-associated genes including *Chchd10, Pcsk1n*, and *Tubb2b*, together with contractile-associated markers such as *Tagln*, and *Actg2*. This mixed expression pattern suggests that State 2 represents a relatively plastic intermediate undergoing fate diversification. In contrast, State 3 showed a more pronounced and restricted contractile program, characterized by *Tagln, Acta2, Actg2*, and *Myl9*. The dot plot and matrix plot summarize the mean expression and cellular prevalence of these biologically informative markers across the sampled generated cells. Accordingly, the three states were annotated as a matrix-associated stromal-like state, a less-committed reprogramming intermediate, and a contractile stromal-like state, respectively. A complementary model-based analysis of the expression distribution *p*(*X*_*g*_ |*c*) (defined in Supplementary Note 5), where *X*_*g*_ denotes the gene-expression variable for canonical gene *g*, showed broadly consistent state-level transcriptional patterns (Supplementary Fig. 5 (b)).

## 3 Discussion

Cell evolution is a dynamic process that can be understood by studying scRNA-Seq time course data. However, effectively modelling the data faces two main challenges. First, the destructive nature of the technology prevents us from following the same cell over time. Second, we cannot observe cells at all timepoints along the trajectory. To tackle this problem, various approaches have been proposed to reconstruct cellular trajectories and track cell evolution from time-course single-cell data. One line of work reconstructs gene expression profiles at unobserved intermediate stages by interpolating between adjacent observed time points, rather than by modeling the underlying continuous-time dynamics [26][8][27][5]. Another major paradigm explicitly models cellular kinetics to reconstruct continuous progression. These methods learn global dynamical systems that govern the evolution of cell-state distributions over time [28][4]. They typically model cellular transitions as continuous velocity fields using neural ordinary differential equations [29] or stochastic differential equations [28][30], with extensions to unbalanced settings that account for cell proliferation and death [31][32][6]. However, they still have limitations. For methods that explicitly models cellular kinetics to reconstruct continuous progression, they primarily learn the direction of the transition from the start to the end, but give less consideration to the distribution of gene expression at each observed timepoint. In methods that are based on interpolation between observed stages without modeling the underlying continuous-time dynamics, even though they consider the distribution of gene expression at each observed timepoint, it is challenging for them to actually capture the dynamic information across all cells along the trajectory. Moreover, all existing methods cannot capture the hidden state of each cell, which is an essential factor to determine gene expression profiles at different timepoints.

Therefore, as shown in Fig. 1, we propose a method that is able to capture the cell state dynamics and gene expression distribution of cells at different timepoints to recover the whole trajectory. We compare our model with three state-of-the-art approaches, scNode[4], ARTEMIS[6] and scDiffusion[5], on the Pancreatic dataset [7] and Waddington-OT dataset [8]. As shown in Fig. 2 and Fig. 3, our model obtains the best interpolation performance to recover the gene expression of cells at the unobserved timepoints. A key feature of our work is that the Latent Cell-State Model not only generates realistic gene expression of each cell at each timepoint, but also provides a corresponding biologically meaningful cell state. As shown in Fig. 4 and Fig. 5, many known gene markers can be identified in each cell state captured by our model to determine its biological meaning, making the cells generated by our model biologically explainable.

Our current work primarily focuses on studying cellular evolution using scRNA-Seq time-course data. To capture a more complete picture of evolutionary dynamics beyond transcriptomics alone, in the future, we will extend our framework to consider cellular evolution across multiple molecular layers, thereby enhancing the versatility and robustness of our model.

## 4 Methods

To make it easy for explanation, we illustrate the details of our model using one sample. Then the likelihood of one cell sample can be written as:

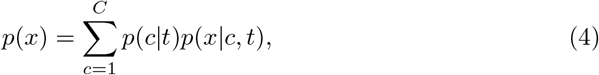

where

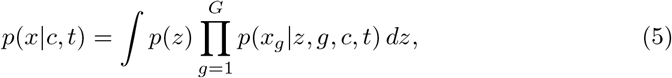

*x* denotes the gene expression profiles for one cell sample, *t* indicates the time when the cell is destroyed to measure the gene expression, *c* denotes the cell state at time *t, z* denotes the latent variable, which captures higher-dimensional features of a cell in a lower dimension, representing the remaining variation within that cell, *g* indicates a gene, and *x*_*g*_ indicates the expression of the gene *g*.

### 4.1 Cell State Modeling

As cell state transitions occur along the trajectory, we assume that one cell starts in some state distribution *π*_*i*_ at time 0, where *i ∈* {1, …, *C*}, and evolves according to a Markov transition matrix *P* shared across all cells along the trajectory, so that at time *t* the distribution is

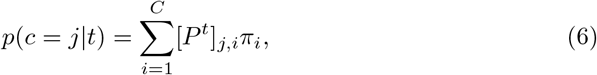

where *P*^*t*^ is *t*^*th*^ power of the Markov transition matrix *P*. *P* can be parametrised using a softmax:

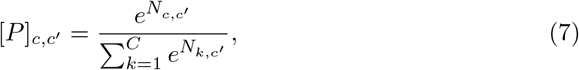

where *c*′ denotes the cell state at time *t*−1 and *N* is a *C* × *C* parameter matrix. To prevent cells from revering to earlier states while still allowing the model to learn transition probabilities, we define *N* to be an upper triangular matrix to prevent transitioning to earlier states:

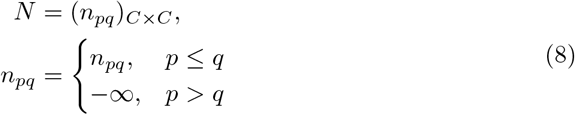

where the upper triangular entries *n*_*pq*_ (*p* ≤ *q*) are learnable parameters updated during training via gradient descent. Setting *n*_*pq*_ = −∞ for *p > q* guarantees *e*^*−∞*^ = 0, so that backward transitions receive zero probability after the softmax, enforcing a strict left-to-right Markov structure. Moreover, the initial distribution *π* is shared across all cells along the trajectory, parametrized by a softmax:

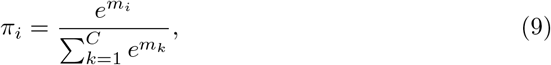

where **m** = (*m*_1_, *m*_2_, …, *m*_*C*_)^*⊤*^ *∈* ℝ^*C*^ is a learnable parameter vector.

### 4.2 Gene Expression Modeling

Then, we need to model the cellular gene expression. In our model, expression counts of each gene in each cell are represented by a negative binomial distribution condition on the current cell state *c*, time *t*, a latent variable *z* and the gene *g*, reflecting the underlying Poisson-gamma mixture model. For each cell, let 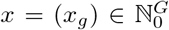 denote the count vector, where *G* is the number of genes in one cell, and *x*_*g*_ represents the count of a gene *g* in this cell. Given the sparsity and overdispersion of gene expression, we assume that *x*_*g*_ follows the negative binomial distribution:

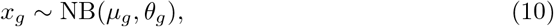

where *µ*_*g*_ *∈* ℝ_*≥*0_ and *θ*_*g*_ *∈* ℝ_*>*0_ are cell-gene-specific mean and inverse dispersion.

The negative binomial distribution can be parameterized by the following probability mass function:

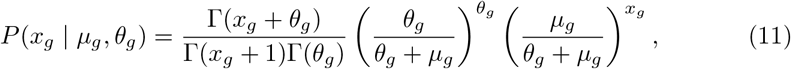

where Γ(·) denotes the Gamma function. The mean and variance of this distribution are given by E[*x*_*g*_] = *µ*_*g*_ and Var 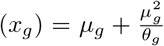, respectively [33]. In contrast to the Poisson distribution, where Var(*x*_*g*_) = *µ*_*g*_, the negative binomial distribution allows for overdispersion. Moreover, *µ*_*g*_ can be further decomposed as *µ*_*g*_ = *ℓ* · *ρ*_*g*_, where *ℓ* denotes the total count of a cell, serving as a scaling factor, and *ρ*_*g*_ represents the relative abundance of the gene [34].

### 4.3 Variational Objective and Formulation

To evaluate the likelihood of one cell shown in Eq. 4, we need to obtain *p*(*x*|*c, t*) illustrated in Eq. 5 (that is equivalent to 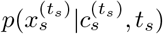 in Eq. 2). As marginalize over the latent variable *z* via the integral 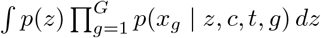 is computationally intractable, a variational approach is used to bound it.

To make it easy for explanation, we dropout *g* from 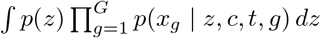 and rewrite it as ∫*p*(*z*)*p*(*x* | *z, c, t*) *dz*. Then, we introduce a variational distribution *q*(*z* | *x, c, t*) following the Gaussian distribution as an approximation to the true posterior *p*(*z* | *x, c, t*), thereby transforming the intractable integration problem into the maximization of a lower bound. Based on the non-negativity of the KL divergence, the lower bound can be derived as follows:

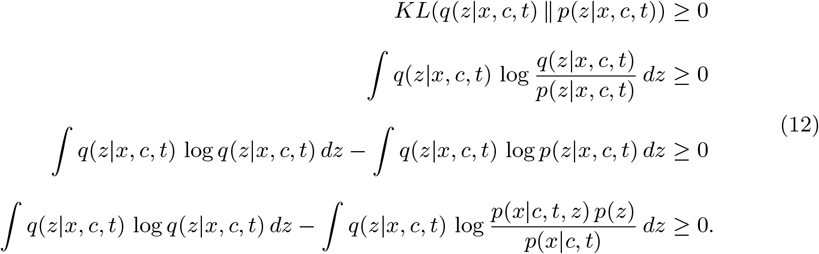

Then the lower bound of the log *p*(*x*|*c, t*) can be obtatined as follows:

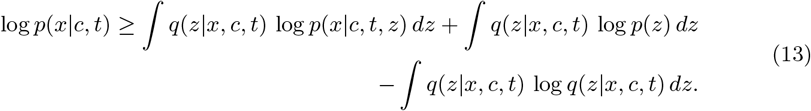

After obtaining the lower bound of log *p*(*x*| *c, t*), we can derive the evidence lower bound (ELBO) of the log-likelihood *p*(*x*) for one cell sample as follows:

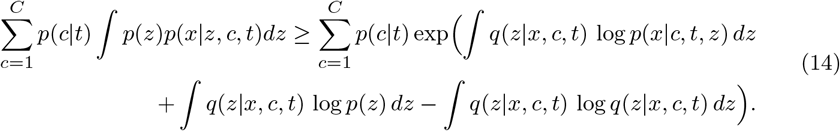

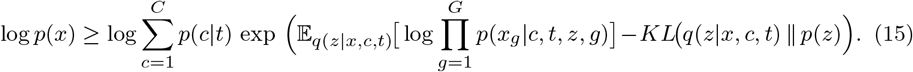

And to obtain the likelihood of all cell samples *p*(X) in Eq. 1, we need to obtain 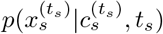 in Eq. 2 that is equivalent to *p*(*x*|*c, t*) in Eq. 5. Based on the lower bound of log *p*(*x*|*c, t*) in Eq. 16, the lower bound of log 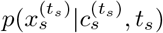 can be represented as:

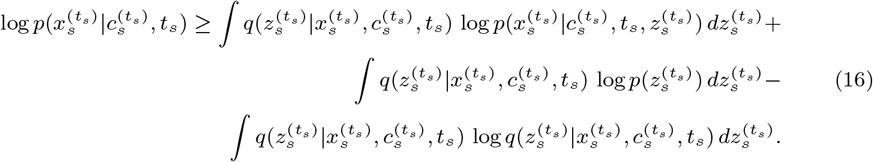

Then we can derive the ELBO of the log-likelihood *p*(X) for all cell samples as follows:

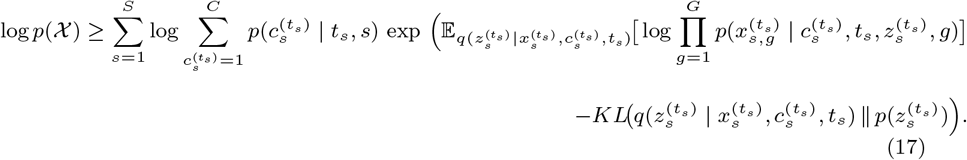

### 4.4 Neural Parameterization and Implementation

The optimization process shown above is carried out within a variational framework. Further details on the implementation and training configurations of the neural networks are provided in Supplementary Notes 1 and 2.

Firstly, we need to parametrize cell state *c*, gene label *g*, and time *t*. The cell state *c* is parametrized by a cell state embedding *e*_*c*_. Since, there is no ordering requirement for the cell state *c*, the cell state embedding *e*_*c*_ can be represented as:

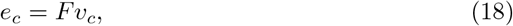

where *v*_*c*_ is a one-hot vector with a 1 at position *c* and 0s elsewhere. 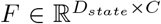 is a learnable matrix, where *C* denotes the number of cell states and *D*_*state*_ is a hyperparameter.

The gene label is parametrized by a gene embedding vector *e*_*g*_. Since there is no ordering requirement for the gene *g*, the gene embedding *e*_*g*_ can be represented as:

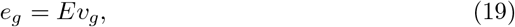

where *v*_*g*_ is a one-hot vector with a 1 at position *g* and 0s elsewhere. 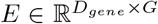 is a learnable matrix, where *G* denotes the number of genes in the given dataset and *D*_*gene*_ is a hyperparameter.

The time *t* can be parametrized by a time embedding *e*_*t*_. To ensure the smoothness of the time *t*, it is represented by:

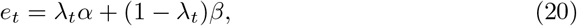

where *α* and *β* are learnable vectors in 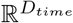 and *D*_*time*_ denotes the dimension of the time embedding *e*_*t*_, which is a hyperparameter. Additionally,

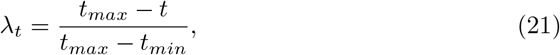

where *t*_max_ and *t*_min_ represent the endpoints of the time interval along the trajectory.

#### 4.4.1 Model Training

During the inference process, the latent variable 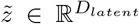 can be obtained as follows:

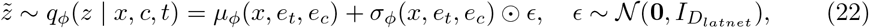

where 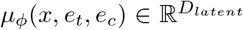 and 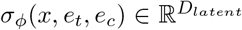 are the mean and standard deviation output by the neural network *ϕ* and 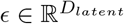 is a random vector sampled from a standard Gaussian, *⊙* denotes element-wise multiplication, and *D*_*latent*_ is the latent dimension.

The reconstruction expression of each gene 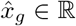 is obtained as follows:

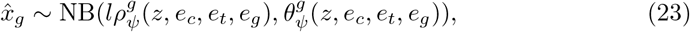

where 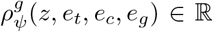 is the relative abundance of the gene, *l* denotes the total count of the cell. 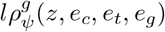 indicates the mean of the negative distribution, and 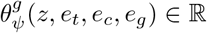 is the inverse dispersion. Both 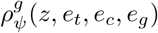 and 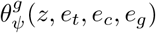 are output by the neural network *ψ*.

Finally, the neural networks *ϕ* and *ψ* are jointly trained, following loss function:

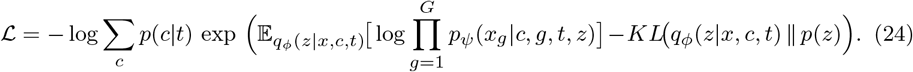

#### 4.4.2 Generative Sampling

Once the model is trained, new cell samples can be generated without the neural network *ϕ*. The latent variable 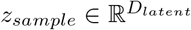 can be sampled as follows:

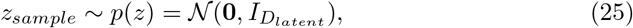

where *D*_*latent*_ is the latent dimension, 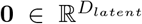 is the zero mean vector and 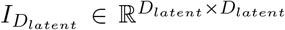 is the identity covariance matrix. Afterwards, the initial cell state is sampled according to the probability of each cell state obtained by Eq. 9 and the cell state *c*_*sampling*_ at current time *t*_*sampling*_ is sampled according to the probability of each cell state obtained by Eq. 6. Then, the embedding of the cell state, the time embedding, the latent variable *z*_*sampling*_, and the embedding of the gene are passed through the neural network *ψ*, where all genes in one cell are iterated and passed through the neural network *ψ* sharing the same time and cell state to obtain gene expression profile for one cell.

### 4.5 Data Preprocessing

To evaluate the quality of the samples generated by our model at unobserved timepoints, we selected two representative datasets: the Pancreatic dataset [7], corresponding to stage 5 of the differentiation protocol, and the Waddington-OT dataset [8], spanning days 0 to 8 under doxycycline (dox) induction.

For the Pancreatic dataset, all cells span from day 0 to day 7 (8 timepoints), therefore, the number of timepoints *T* is set to 8. When training models in the Pancreatic dataset to evaluate interpolation performance, days 3 and 6 are set as held-out days. Therefore, the Pancreatic training dataset (cells can be observed in practice) including all cells except samples on days 3 and 6 is used to train the model to evaluate the interpolation performance, where S is 51274 (See Supplementary Tab. 2 for details). Then, we select 2000 samples on each held-out day to construct the Pancreatic test dataset (cells that cannot be observed in practice).

For the Waddington-OT dataset, all cells span from day 0 to day 8 with a half-day interval (17 timepoints). Therefore, when training our model in the Waddington-OT dataset, the number of timepoints *T* is set to 17. Days 3.5 and 4 are set as held-out timepoints to evaluate the interpolation performance of all models. Therefore, the Waddington-OT training dataset (cells can be observed in practice) including all cells except samples on days 3.5 and 4 is used to train the model to evaluate the interpolation performance, where S is 73693 (See Supplementary Tab. 1 for details). Then, we select 1000 samples on each held-out day to construct the Waddington-OT test dataset (cells that cannot be observed in practice).

Then three main steps are implemented on each dataset: 1) 2000 highly variable genes (HVGs) are identified from the training datasets, using method proposed in [35]; The genes selected from the training datasets are then used to subset the corresponding test datasets, so the number of genes *G* is fixed at 2000 across all datasets; 2) For comparison methods requiring normalized expression values, counts are normalized by depth scaling using the median count of each dataset as the target count, ensuring that each cell has a consistent total count across all genes. 3) For comparison methods requiring normalized expression values, log1p normalization, i.e., log transformation after adding one, is applied to stabilize the variance. For the datasets used to train our model, we only apply the first step, as our model takes count data as input. However, for the datasets used to train the comparison methods, we apply all three steps, since these methods require normalized data as input.

## Acknowledgement

We express our sincere gratitude to Dr. Arthur Radley from the Francis Crick Institute for insightful discussions on biological interpretation.

## Data availability

The source data supporting the findings of this study are available upon reasonable request during the peer-review process.

## Code availability

The custom computer code and evaluation algorithms central to the main claims of this paper are available to the editors and reviewers upon request for peer-review purposes.

## Supplementary information

Additional details are provided in the Supplementary Information file.

## Supplementary Information

**Supplementary Table 1.** Number of cells per timepoint in training dataset from Pancreatic dataset.

| **Timepoint (Day)** | **Number of Cells** |
|---|---|
| 0 | 6190 |
| 1 | 6177 |
| 2 | 3562 |
| 4 | 6480 |
| 5 | 6404 |
| 7 | 3971 |

**Supplementary Table 2.** Number of cells per timepoint in training dataset from Waddington-OT dataset.

| **Timepoint (Day)** | **Number of Cells** |
|---|---|
| 0 | 3911 |
| 0.5 | 2939 |
| 1 | 3198 |
| 1.5 | 1687 |
| 2 | 8416 |
| 2.5 | 6024 |
| 3 | 5739 |
| 4.5 | 6226 |
| 5 | 6241 |
| 5.5 | 5759 |
| 6 | 7368 |
| 6.5 | 2737 |
| 7 | 5674 |
| 7.5 | 4389 |
| 8 | 3385 |

**Supplementary Figure 1.**
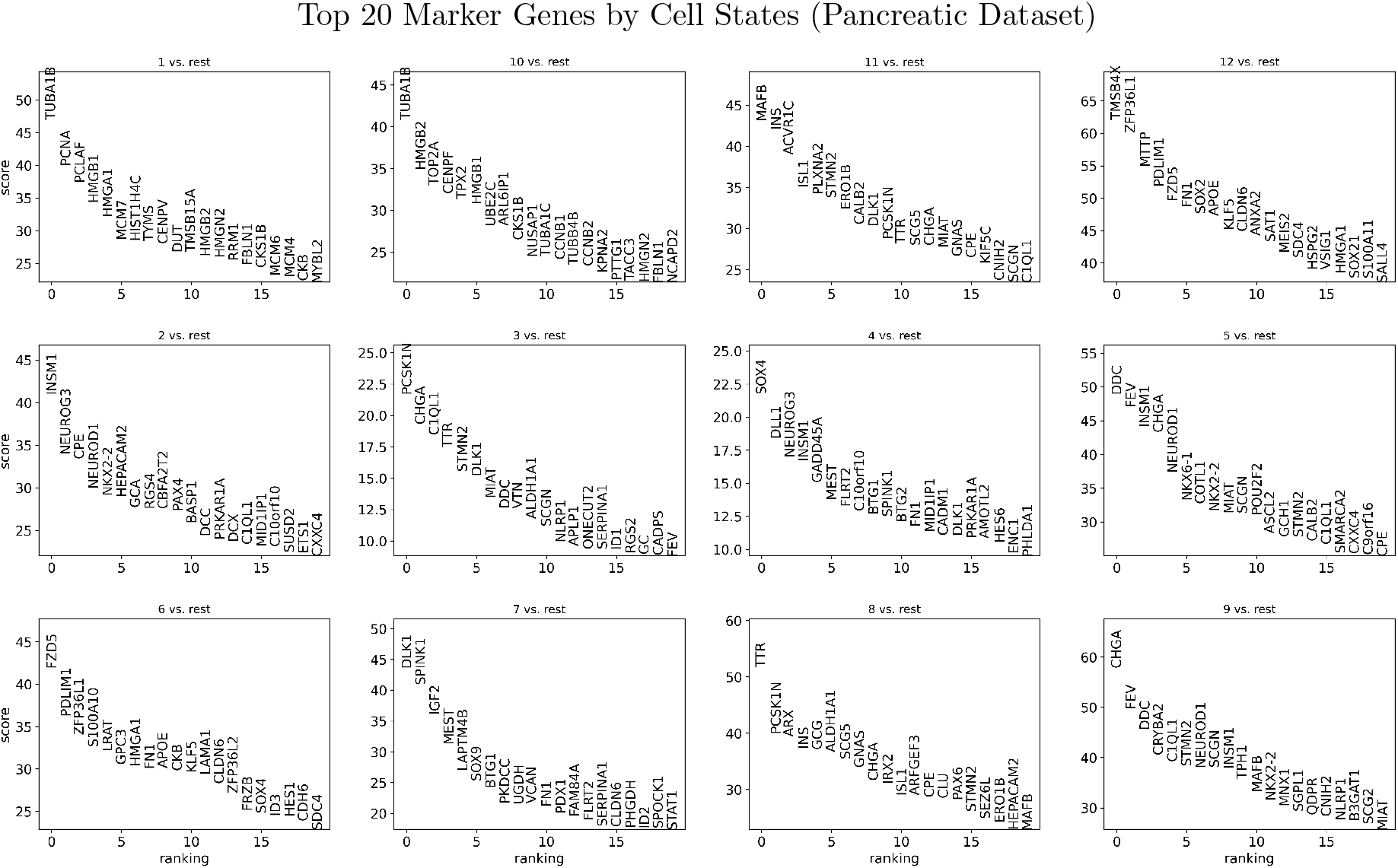
Top 20 marker genes identified for each cell state given by our model in the Pancreatic dataset.

**Supplementary Figure 2.**
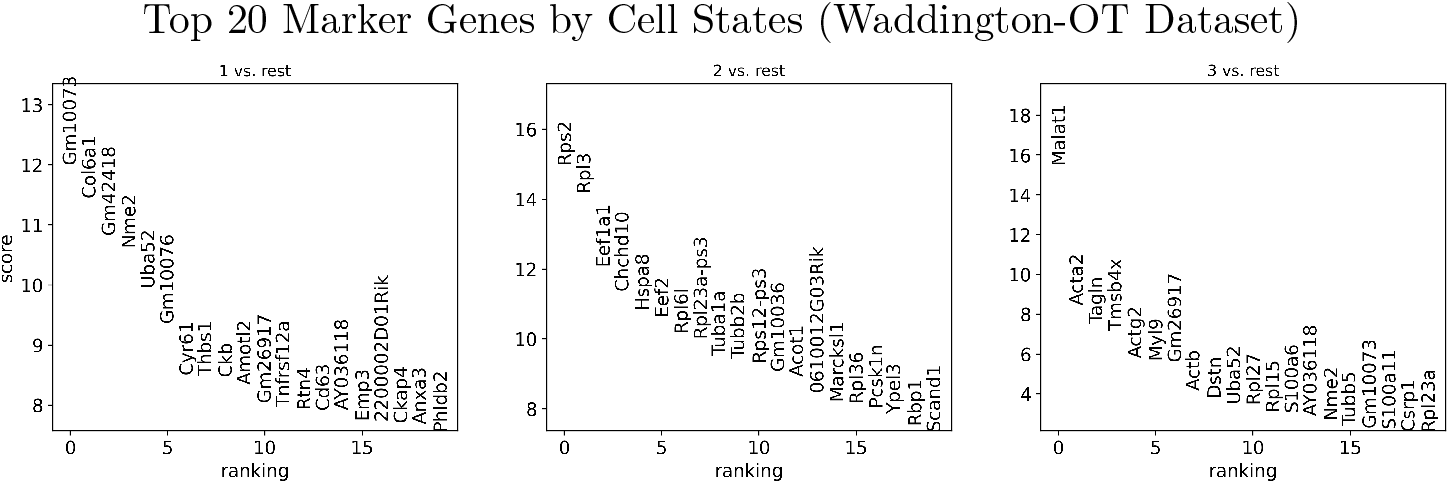
Top 20 marker genes identified for each cell state given by our model in the Waddington-OT dataset.

### Supplementary Note 1. Implementation Details

We assume that the latent variable, the gene type, the time, and the cell state contribute equally to the expression of each gene. Then, we assign the same dimension to each of them. The dimension of the latent variable 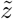 for training and *z*_*sample*_ for sampling, i.e., *D*_*latent*_, the *D*_*time*_ dimension of time *t*, i.e., *e*_*t*_, the *D*_*state*_ dimension of the cell state *c*, i.e., *e*_*c*_ and the *D*_*gene*_ dimension of the gene type *g*, i.e., *e*_*g*_ are set according to the ablation study for the relevant datasets and tasks. Details on how their optimal dimensions are selected for the Pancreatic and Waddington-OT datasets are provided in Supplementary Note 2.

Then, the input of the neural network *ϕ* of our model is a vector concatenated by *e*_*c*_, *x*, and *e*_*t*_. It is then inputted into a fully connected layer, followed by a normalization layer and a ReLU layer to reduce the dimension to 512. Another fully connected layer, followed by a normalization layer and a ReLU layer, is added to further reduce the dimension to 256. Afterward, two fully connected layers are added to obtain the vector with dimension *D*_*latent*_. For the neural network *ψ*, the input is a vector concatenated by *e*_*g*_, *e*_*c*_, *e*_*z*_ and *e*_*t*_. Afterward, it is input into one fully connected layer, followed by a normalization layer and a ReLU layer to reduce the dimension to 512. Then another fully connected layer, followed by a normalization layer and a ReLU layer, is added to further reduce the dimension to 256. Finally, a fully connected layer is added to reduce the dimension to 2. The proposed model is trained on Pytorch 2.5.1 and CUDA 12.1. They are optimized using the Adam optimizer with a learning rate of 10^*−*3^ and a maximum of 100 iterations.

### Supplementary Note 2. Hyperparameters Selection

To determine the optimal values of several key hyperparameters including *C, D*_*state*_, *D*_*time*_, *D*_*gene*_, and *D*_*latent*_ in our neural network, we conducted ablation studies. For both Waddington-OT and Pancreatic datasets, the validation dataset for hyperparameter selection is constructed from the training dataset described in Section 4.5 by holding out 300 samples from each day for evaluation, while the remaining samples are used to train the model.

- *C* **(Number of Cell States)**

**Supplementary Figure 3.**
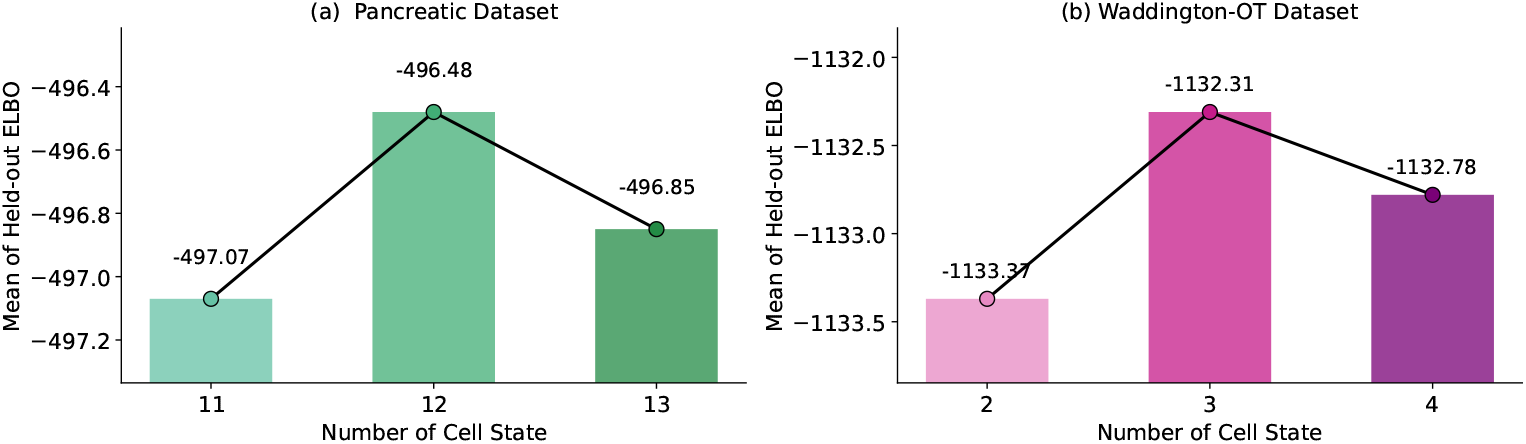
Number of Cell States: (a) The mean ELBO of all cell samples in the Pancreatic validation dataset; (b) The mean ELBO of all cell samples in the Waddington-OT validation dataset.

An ablation study has been conducted to determine the optimal number of cell states across different datasets. Performance is determined by the mean ELBO of the cells in the validation datasets. It is shown that the model with 3 cell states achieves the best performance on Waddington-OT dataset, shown in Supplementary Fig. 3(b). For the Pancreatic dataset, the model with 12 cell states can gain the highest performance, shown in Supplementary Fig. 3(a). Therefore, when using the Waddington-OT dataset to train our model and compare its interpolation performance with other methods, the number of cell states *C* is set to 3. And we set the number of cell states *C* to 12 when using the Pancreatic dataset to train our model, in order to compare its interpolation performance with other methods.

- *D*_*latent*_, *D*_*gene*_, *D*_*time*_, *D*_*state*_ **(Dimension of Latent Variable, Gene Type, Time, Cell State)**

We assume that the latent variable, the gene type, the time, and the cell state contribute equally to the expression of each gene, therefore, we set the same dimension to each of them and conduct the ablation study. The ablation study has been conducted to determine the optimal number of dimensions for time, cell state, gene type, and latent variable. Performance is determined by the mean ELBO of the cells in the validation datasets. The model with 256 dimensions for all variables is shown to achieve the best performance in both Pancreatic and Waddington-OT datasets, as demonstrated in Supplementary Fig. 4(a) and (b). Therefore, we set the dimensions of the latent variable *D*_*latent*_, the gene type *D*_*gene*_, the time *D*_*time*_ and the cell state *D*_*state*_ to 256, when training our model to compare its performance with other methods.

**Supplementary Figure 4.**
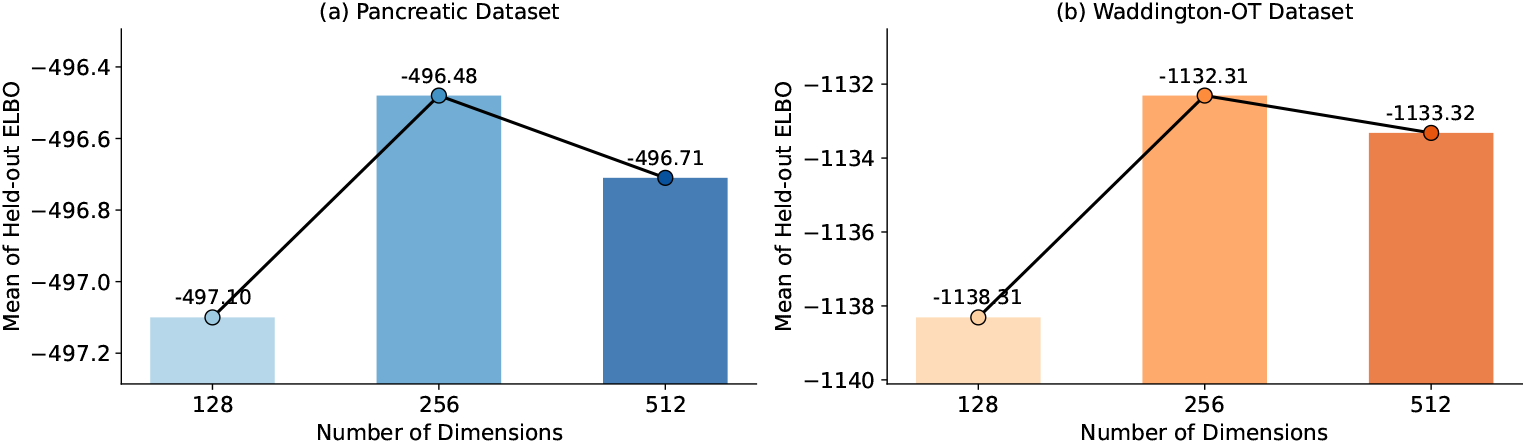
Number of Dimensions: (a)The mean ELBO of all cell samples in Pancreatic validation dataset; (b) The mean ELBO of all cell samples in validation Waddington-OT validation dataset

### Supplementary Note 3. Evaluation Matrices

#### Maximum Mean Discrepancy (MMD)

Maximum Mean Discrepancy (MMD) is used to quantify the discrepancy between the real and generated data distributions, which enables direct comparison of distributions from samples without requiring density estimation and explicit geometric assumptions on the data space. MMD is measured in a 50-dimensional principal component analysis (PCA) space [12]. PCA is fitted on the real samples only, and generated samples are embedded into this fixed space using the principal component loadings learned from the real data. Since our model generates count data, while other comparison methods generate normalized expression values, we preprocess the generated and real samples consistently before PCA. For each held-out time point, real samples are normalized by depth scaling, where the scaling factor is defined as the median total count of all real samples from the corresponding day in the test dataset. A log1p transformation is then applied to stabilize the variance. Generated samples are processed using the same normalization and log1p transformation pipeline before projection into the real-data PCA space.

The MMD is formulated as follows:

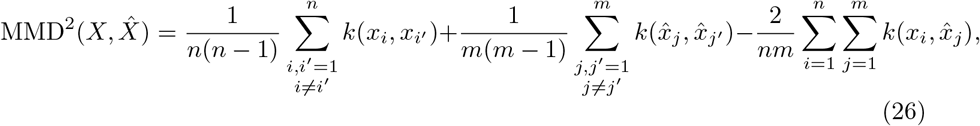

where 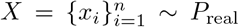 denotes samples from real distribution and *n* denotes the number of samples in *X*. 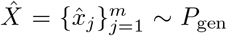 indicates samples from generated distribution and *m* denotes the number of samples in 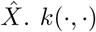 is a Gaussian kernel defined as:

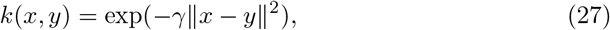

where *∥* · *∥* denotes the Euclidean norm. And, the kernel parameter *γ* is determined using the median heuristic as follows:

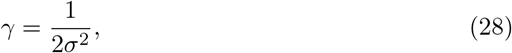

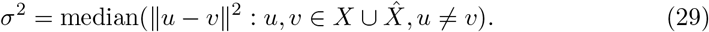

#### UMAP Visualization of Real and Generated Samples

UMAP visualization is used to project each 2000-dimensional gene expression sample into a two-dimensional space for qualitative assessment. We follow the same method-specific preprocessing strategy as in the MMD evaluation. For our count-generating model, samples are normalized and log1p-transformed, whereas samples from comparison methods that generate normalized expression values are used directly. A balanced real reference set is constructed by sampling an equal number of real cells from each observed date, matching the number of samples per held-out date in the test dataset. UMAP is fitted exclusively on this real reference set, and generated samples are subsequently projected onto the fixed UMAP embedding learned from the real data using the ingest function in Scanpy. The resulting two-dimensional representations are used to qualitatively assess how well generated samples align with real samples at each held-out time point.

### Supplementary Note 4. Cell State Annotation via Scanpy

#### UMAP Visualization of Generated Cell States

Gene expression counts were normalized to the median total count per cell and subsequently log1p-transformed. UMAP visualization of generated samples colored by their corresponding sampled cell states. Generated cells were projected onto the same fixed UMAP embedding derived from real reference cells using Scanpy’s `ingest` function [11]. Each point represents a generated cell, colored according to its corresponding sampled cell state label. Distinct colors indicate different sampled cell states.

#### Cluster-Specific Marker Genes Identification

Gene expression counts were normalized to the median total count per cell and subsequently log1p-transformed. Cluster-specific genes were identified using the Wilcoxon rank-sum test implemented in `sc.tl.rank_genes_groups` (Scanpy) [11], comparing each cluster against all remaining cells (one-versus-rest approach). P-values were adjusted for multiple testing using the Benjamini-Hochberg procedure. Genes were ranked in descending order of their Wilcoxon test scores, and the top 20 ranked genes for each cluster were visualized using `sc.tl.rank_genes_groups` (Scanpy) [11].

#### Hierarchical Clustering of Cell States

Gene expression counts were normalized to the median total count per cell, followed by log1p-transformation. Hierarchical clustering of cell states was performed using `sc.tl.dendrogram` in Scanpy [11], based on the first 50 principal components (PCs) derived from the normalized and log-transformed gene expression data. Pairwise Pearson correlations were calculated between the cluster-level mean PCA representations, followed by complete-linkage hierarchical clustering. The resulting dendrogram was visualized using `sc.pl.dendrogram`.

#### Canonical Marker Gene Expression Visualization

Following the identification of cluster-specific genes, canonical marker genes from the reference were cross-referenced with the top 20 Wilcoxon-ranked genes of each cell state. Only canonical marker genes present in these top-ranked gene lists were selected for visualization using `sc.pl.dotplot` and `sc.pl.matrixplot` in Scanpy [11].

For each gene *g* in cell state *c*, the mean expression level was calculated as

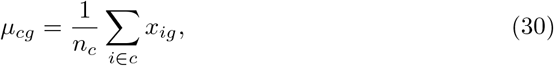

where *c* denotes set of cells in state *c, n*_*c*_ is the number of cells in that state, and *x*_*ig*_ represents the normalized and log1p-transformed expression value of gene *g* in cell *i*.

Mean expression levels were independently scaled to the range [0, 1] for each gene across cell states using min–max scaling:

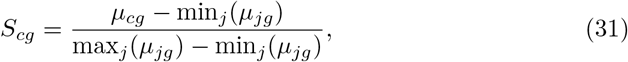

where *µ*_*cg*_ represents the mean expression level of gene *g* in cell state *c*, and *j* indexes all cell states. The terms min_*j*_(*µ*_*jg*_) and max_*j*_(*µ*_*jg*_) denote the minimum and maximum mean expression levels of gene *g* across all cell states, respectively. In the dot plot, dot color represents the scaled mean expression *S*_*cg*_, whereas dot size *P*_*cg*_ reflects the fraction of cells with non-zero expression of the corresponding gene:

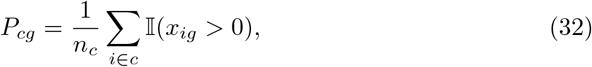

where I(·) is the indicator function. In the matrix plot, the scaled mean expression values were represented using a color gradient.

### Supplementary Note 5. Refined Cell State Analysis

To further evaluate the biological relevance of the cell states learned by our model, we estimate the conditional distribution *p*(*X*_*g*_ | *c*) of expression levels *X*_*g*_ for canonical driver genes *g* of the targeted biological process across cell states *c ∈ C*, where *p*(*X*_*g*_| *c*) is defined as:

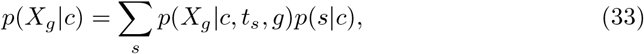

We assume that *p*(*s* | *c*) *∝ p*(*c* | *s, t*_*s*_), where *s* denotes a single sample among all generated samples and *t*_*s*_ denotes the timestamp of sample *s*. Consequently, *p*(*s* | *c*) can be defined as:

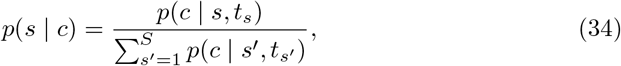

For clear visual inspection, we present the expected expression level E[*X*_*g*_ |*c*] using a heatmap structured as follows, where

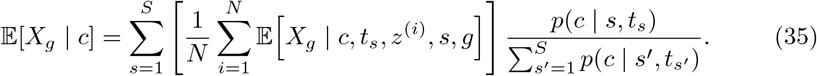

The value *v*_*cg*_, encoded by the color intensity of each unit in the heatmap, is calculated as follows:

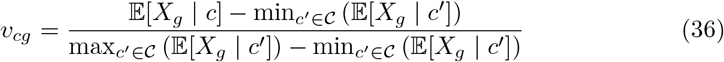

We plot the heatmaps defined above for the canonical genes in the Pancreatic dataset [7] and Waddington-OT dataset [8], as shown in Supplementary Fig. 5(a) and Supplementary Fig. 5(b), respectively. They exhibit the similar patterns as the dot plots and matrix plot shown in Fig. 4(c) and (d) and Fig. 5(c) and (d), which are generated using the plotting approach defined in [11].

**Supplementary Figure 5.**
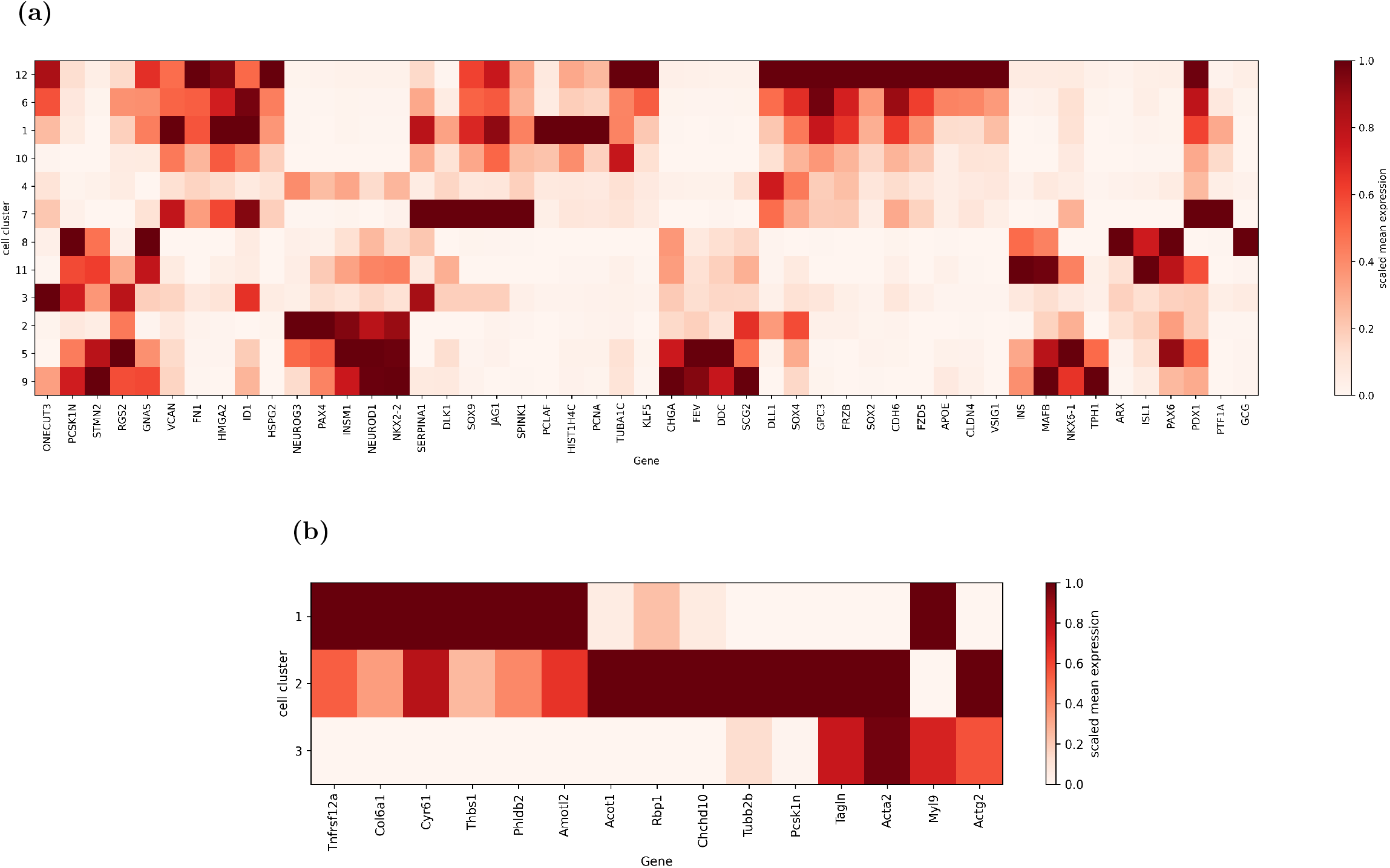
Heatmaps obtained using the method described in Supplementary Note 5. (a) Pancreatic dataset; (b) Waddington-OT dataset.

## Footnotes

1 This is analogous to non-linearly projecting the high-dimensional gene expression vector to a lower-dimensional space shared by all cell samples, cell-states, and times.

